# Liquid-like condensates mediate competition between actin branching and bundling

**DOI:** 10.1101/2023.06.23.546267

**Authors:** Kristin Graham, Aravind Chandrasekaran, Liping Wang, Noel Yang, Eileen M. Lafer, Padmini Rangamani, Jeanne C. Stachowiak

## Abstract

Cellular remodeling of actin networks underlies cell motility during key morphological events, from embryogenesis to metastasis. In these transformations there is an inherent competition between actin branching and bundling, because steric clashes among branches create a mechanical barrier to bundling. Recently, liquid-like condensates consisting purely of proteins involved in either branching or bundling of the cytoskeleton have been found to catalyze their respective functions. Yet in the cell, proteins that drive branching and bundling are present simultaneously. In this complex environment, which factors determine whether a condensate drives filaments to branch versus becoming bundled? To answer this question, we added the branched actin nucleator, Arp2/3, to condensates composed of VASP, an actin bundling protein. At low actin to VASP ratios, branching activity, mediated by Arp2/3, robustly inhibited VASP-mediated bundling of filaments, in agreement with agent-based simulations. In contrast, as the actin to VASP ratio increased, addition of Arp2/3 led to formation of aster-shaped structures, in which bundled filaments emerged from a branched actin core, analogous to filopodia emerging from a branched lamellipodial network. These results demonstrate that multi-component, liquid-like condensates can modulate the inherent competition between bundled and branched actin morphologies, leading to organized, higher-order structures, similar to those found in motile cells.

**SIGNIFICANCE STATEMENT:** Reorganization of actin filaments allows cells to migrate, which is required for embryonic development, wound healing, and cancer metastasis. During migration, the leading-edge of the cell consists of needle-like protrusions of bundled actin, which emanate from a sheet of branched actin. Given that the proteins responsible for both architectures are present simultaneously, what determines whether actin filaments will be branched or bundled? Here we show that liquid-like condensates, composed of both branching and bundling proteins, can mediate the inherent competition between these fundamentally different ways of organizing actin networks. This work demonstrates that by tuning the composition of condensates, we can recapitulate the transition from branched to bundled networks, a key step in cell migration.

## INTRODUCTION

The cellular cytoskeleton is organized into numerous higher-order structures that achieve diverse physiological functions. Specifically, the actin cytoskeleton can be organized into parallel or antiparallel bundles, branched networks, or crosslinked networks, each conferring different mechanical properties that are optimized for specific cellular processes (1–3). Cell motility relies on the combined efforts of these different morphologies to enable propulsion of the cell’s leading edge, central contraction, and subsequent retraction of the cell’s trailing edge. The leading edge comprises both parallel actin bundles, found in finger-like filopodial protrusions, and branched actin networks, found in the sheet-like lamellipodium (4). Assembly of branched actin networks requires the branch nucleator, Arp2/3, which, upon activation by Wiskott-Aldrich syndrome protein (WASP), binds to the sides of mother actin filaments, nucleating a branch (5,6). Interestingly, assembly of parallel bundles of actin filaments, found in filopodia, is thought to originate from an underlying branched actin network. Specifically, processive actin polymerases, such as formins or Ena/VASP proteins, are thought to catalyze the convergent elongation of linear filaments that originate from the branched actin network of the lamellipodium (7,8). With assistance from crosslinkers like fascin, these parallel bundles can become rigid enough to bend the plasma membrane, leading to protrusions (9).

We have recently shown that the processive actin polymerase, VASP, forms liquid-like droplets that polymerize actin filaments and organize them into parallel bundles, similar to filopodia (10). This process is mediated by competition between the rigidity of the filaments and the surface tension of the droplet, along with VASP’s ability to crosslink filaments. Droplet-mediated bundling of actin occurs through a series of morphological transitions and symmetry breaking events. Specifically, as actin polymerizes, it forms a three-dimensional shell within the droplet, which collapses into a two-dimensional ring as actin polymerization continues. As the ring accumulates more actin and thickens, its increasing bending rigidity eventually overcomes the surface tension of the droplet, deforming the droplet into a disc. Deformation into a disc is concomitant with droplet elongation into higher aspect ratio structures, which eventually form rods. The resulting rods appear to be composed of parallel bundles of actin filaments, consistent with VASP’s cellular function in mediating parallel bundling of linear actin filaments.

Interestingly, recent work from other groups has shown that liquid-like condensates of TPX2 are capable of nucleating microtubule branches (11). Taken together, these studies suggest that liquid-like condensates can be harnessed to facilitate the two major organizing mechanisms within filament networks: bundling and branching. Given the potential for both competition and cooperation between these mechanisms, we wondered how condensates might modulate network morphology when the protein machinery required for bundling and branching are present simultaneously. Specifically, because VASP has been found to drive actin polymerization in plasma membrane clusters in the lamellipodia, as well as in the tips of filopodia, we reasoned that VASP condensates provide an ideal platform to investigate both branching and bundling behaviors (7,12–14).

How might the introduction of a branched actin network impact the bundling activity of the VASP droplets? Here we show that Arp2/3 can be added to VASP droplets to create branched actin networks within the droplets. As expected, droplets that contain a branched network are less likely to deform into linear structures. Specifically, we found that droplets with more Arp2/3, and therefore more branched actin, are arrested at an initial step in the bundling process. Agent-based modeling of filament dynamics suggests that this effect results from Arp2/3’s ability to reduce the effective length of filaments, making it more difficult for them to become bundled. However, as the actin concentration increased, filament networks transitioned from a branched to a bundled state, deforming droplets into aster-shaped structures consisting of a central hub from which thin, actin-filled spokes emanated. Interestingly, Arp2/3 segregated spontaneously to the hubs of these asters and was simultaneously depleted from the spokes, demonstrating actin-mediated partitioning of protein components within droplets. This work illustrates how liquid-like condensates of actin interacting proteins could modulate the competing processes of actin branching and bundling, critical steps in cytoskeletal dynamics throughout the cell.

## RESULTS

### Arp2/3 and VCA participate in phase separation of liquid-like VASP droplets

To investigate the impact of the branched actin nucleator, Arp2/3, on droplet-mediated bundling of actin filaments, we began by incorporating Arp2/3 into droplets composed of VASP. Specifically, we formed droplets of 20 μM VASP (monomer concentration; 5 μM of VASP tetramer) in the presence of Arp2/3, and its activator, the VCA domain of N-WASP. Droplets are formed by mixing VASP and associated proteins with the crowding agent PEG8000, to a final concentration of 3% (w/v). VASP is a native tetramer (15,16). However, we hereafter refer to the monomeric concentration of VASP so that the actin-to-VASP ratios more intuitively reflect the ratio between actin and actin binding sites on VASP. To visualize whether VASP droplets were capable of localizing Arp2/3 and VCA, we labeled Arp2/3 with maleimide Atto-594 (17), and labeled VCA with maleimide Atto-488 (5). Both proteins were enriched in VASP droplets (Figure 1a). While VASP has been found to bind the proline-rich domain of some WASP-family proteins, there are no known interactions of VASP with the WASP VCA domain (18–20). Similarly, Arp2/3 lacks known VASP-interaction motifs. Therefore, enrichment of these proteins in VASP droplets is likely driven by nonspecific or weak binding interactions, which can appear amplified in concentrated droplet systems (21). While addition of Arp2/3 had little effect on VASP droplets, addition of the VCA fragment resulted in VASP droplets with a distribution of larger diameters in comparison to droplets consisting of VASP alone (Figure 1b,c). This effect, which suggests that VCA reinforces condensation of VASP, could result from non-specific interactions between acidic residues in the VCA fragment with basic residues in VASP (15,22–25). Furthermore, the seemingly larger impact of VCA on VASP droplets compared to Arp2/3 could be attributed to the excess VCA relative to Arp2/3, which we used to mimic previous in vitro actin branching assays (26–30). Importantly, when VASP droplets formed in the presence of Arp2/3 and VCA, they remained liquid-like, fusing rapidly upon contact (Figure 1d). Composite droplets of VASP, Arp2/3 and VCA also exhibited recovery after photobleaching that was comparable to that of droplets consisting of VASP alone (Figure 1e,f). Composite droplets exhibited a slightly slower recovery rate, with a t_1/2_ of 7.24±1.06 s, compared to a t_1/2_ of 4.51±0.58 s for VASP-only droplets, suggesting that the presence of VCA and Arp2/3 may increase the degree of protein-protein interaction within droplets. Overall, these data indicate that the presence of Arp2/3 and VCA does not significantly alter the liquid-like nature of VASP droplets.

**Figure 1.**
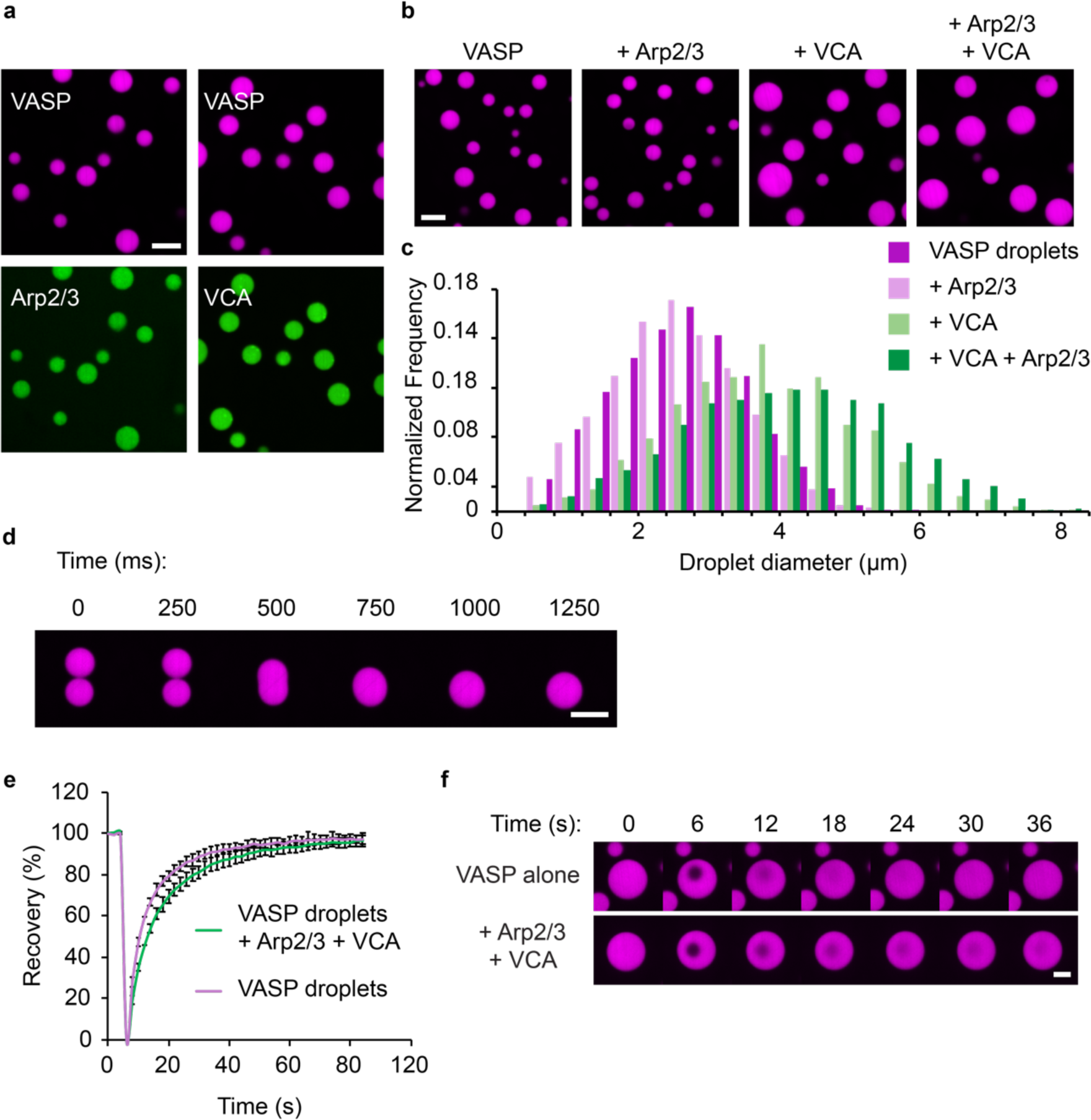
Arp2/3 and VCA participate in phase separation of liquid-like VASP droplets. (a) Enrichment of 1 μM Arp2/3 (labeled with Atto-594, shown in green) or 1 μM VCA (labeled with Atto-488, shown in green) to 20 μM VASP droplets. Droplets are formed in “droplet buffer”: 50 mM Tris pH 7.4, 150 mM NaCl, 3% (w/v) PEG8000. Scale bar 5 μm. (b) Representative images of droplets formed from a 15 μM solution of VASP with either 150 nM Arp2/3, 6 μM VCA, or both 150 nM Arp2/3 and 6 μM VCA. Scale bar 5 μm. (c) Histogram of droplet sizes quantified from (b). Data from n=3 replicates. (d) Composite droplets formed of 15 μM VASP, 150 nM Arp2/3, and 6 μM VCA still remain liquid-like, as exhibited by rapid fusion events. Scale bar 5 μm. (e) Composite droplets formed of 15 μM VASP, 150 nM Arp2/3, and 6 μM VCA display rapid recovery after photobleaching, and the fraction of mobile VASP is nearly equivalent to droplets lacking Arp2/3 and VCA. Curves show the average VASP recovery profile with error bars representing SD for n=11 droplets for each condition. (f) Representative time series of FRAP of either VASP only droplets, or composite droplets as quantified in (e). Scale bar 2 μm.

### Addition of Arp2/3 to VASP droplets inhibits droplet deformation and actin bundling

We next sought to test how branched actin nucleators might alter droplet-mediated bundling of actin filaments. We added 2 μM of actin monomers (labeled with maleimide Atto-488) to pre-formed VASP droplets (15 μM, labeled with maleimide AlexaFluor 647), with increasing concentrations of Arp2/3 and VCA. For each tested concentration of Arp2/3, we maintained the Arp2/3 to VCA ratio constant at 1:40, which is within the range used in literature (26–30). After allowing the actin to polymerize for 15 minutes, after which minimal changes to droplet phenotype are observed (10), the droplets were imaged. As expected from previous work (10), VASP droplets lacking Arp2/3 and VCA deformed into rod-like shapes upon actin polymerization (Figure 2a). However, as the fraction of Arp2/3 and VCA increased, the droplets became more spherical, and their overall aspect ratios decreased (Figure 2a,b). Importantly, actin still polymerized into filaments in the presence of Arp2/3 and VCA, as confirmed with phalloidin staining (Figure 2c). Notably, phalloidin-stained actin frequently arranged into rings at the inner surfaces of droplets, as observed previously (10).

**Figure 2.**
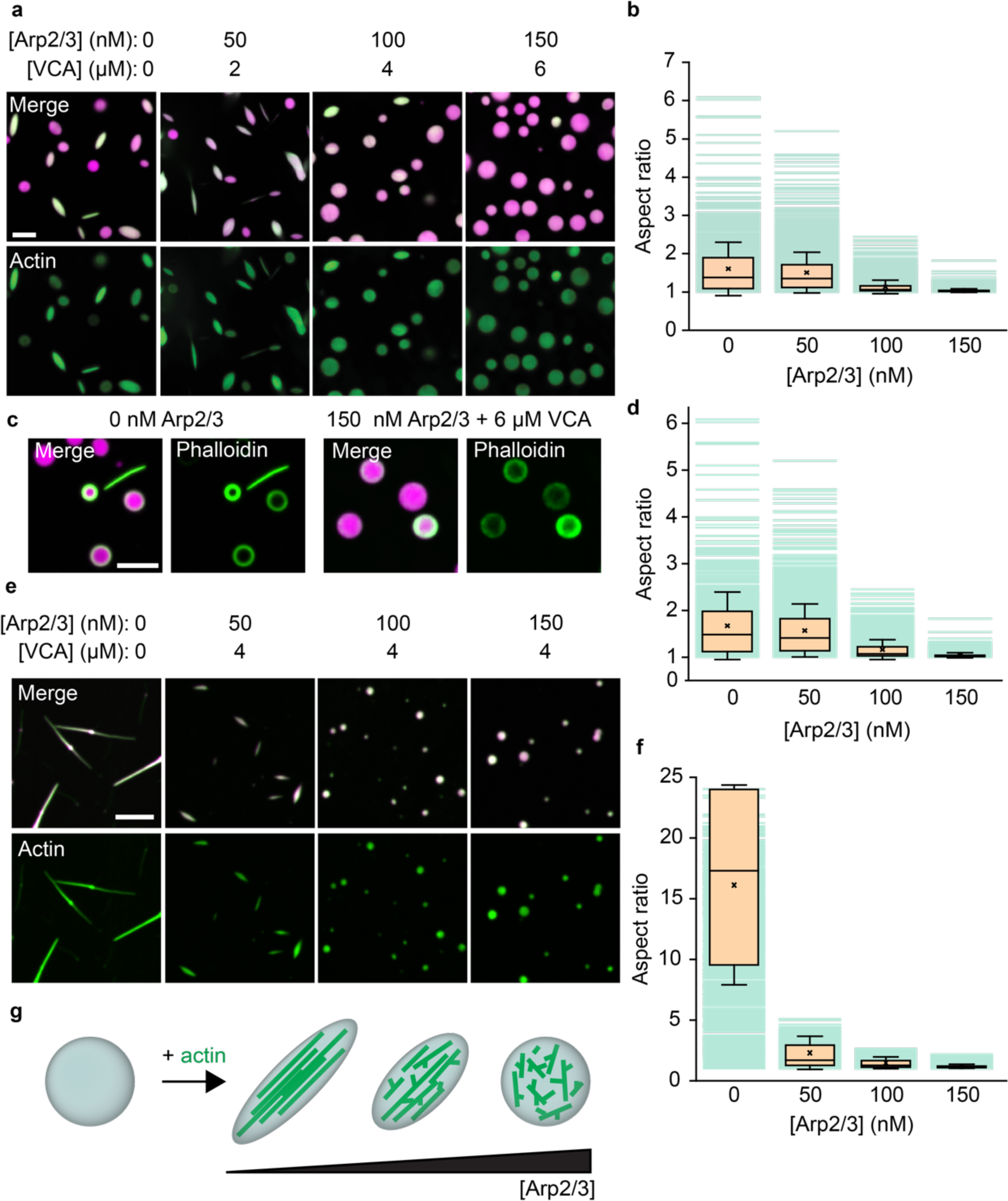
Addition of Arp2/3 to VASP droplets inhibits droplet deformation and actin bundling. (a) Representative images of droplets formed from 15 μM VASP and 2 μM actin, with increasing concentrations of Arp2/3 and VCA. Scale bar 5 μm. (b) Quantification of aspect ratios of droplets in (a). Box represents interquartile range (IQR) with median as a bisecting line, mean as an ‘x’, and whiskers, which represent SD. Each background line represents one data point. Data are from 3 replicates. For 0 nM Arp2/3, n=755 droplets; for 50 nM Arp2/3, n=1391; for 100 nM Arp2/3, n= 2123; for 150 nM Arp2/3, n=2286. (c) Representative images of phalloidin staining of 2 μM actin polymerized by either 15 μM VASP-only droplets, or 15 μM VASP droplets with 150 nM Arp2/3 and 6 μM VCA. Scale bar 5 μm. (d) Quantification of aspect ratios of droplets in (b), filtered for droplets with a minor axis < 3 μm. (e) Representative images of droplets formed from 5 μM VASP, exposed to 0.67 μM actin, with 4 μM VCA and increasing concentrations of Arp2/3. Scale bar 5 μm. (f) Quantification of droplet aspect ratio as a function of Arp2/3 concentration as seen in (e). Box represents IQR with median as a bisecting line, mean as an ‘x’, and whiskers, which represent SD. Data from 3 replicates. For 0 nM Arp2/3, n=656; for 50 nM Arp2/3, n=1116; for 100 nM Arp2/3, n=1239; for 150 nM Arp2/3, n=1006. (g) Cartoon depicting sphere to rod transition driven by the ability of VASP droplets to bundle actin filaments, a process which is inhibited by the addition of Arp2/3 and VCA.

Smaller droplets are more likely to deform at a given actin concentration because growing actin filaments have to bend more tightly to fit inside them (10). Therefore, we asked whether the slight increase in droplet size owing to inclusion of VCA and Arp2/3 (Figure 1c) could explain the reduction in droplet aspect ratio in Figure 2a,b. To answer this question, we reanalyzed the data in Figure 2a,b for droplets with a minor axis below 3 μm, the average size of droplets consisting of VASP alone. Similar to the full population, we observed a substantial reduction in aspect ratio with increasing Arp2/3 concentration for droplets below this threshold (Figure 2d). This result confirms that increased droplet size cannot explain the reduction in droplet deformation in the presence of VCA and Arp2/3. Collectively, these data demonstrate that inclusion of Arp2/3 and VCA in VASP droplets inhibits the deformation of droplets by polymerization of actin.

The experiments in Figure 2a-d examined relatively large VASP droplets for which actin polymerization drives relatively small changes in droplet aspect ratio. To determine whether inclusion of Arp2/3 and VCA can also inhibit larger, actin-driven deformations, we titrated the same concentrations of Arp2/3 in smaller droplets formed at a VASP concentration of 5 μM, rather than 15 μM. Actin was added to these droplets at a concentration of 0.67 μM to maintain the same actin to VASP ratio of 1:7.5 (Figure 2e). The concentration of VCA was 4 μM. In the absence of Arp2/3, actin polymerization within these smaller droplets resulted in rod-like structures with aspect ratios exceeding 20 (Figure 2e, left). However, addition of as little as 50 nM of Arp2/3 resulted in a dramatic decrease in aspect ratio (Figure 2e,f). In particular, droplets of aspect ratio substantially above 1 largely disappeared from the population upon introduction of even 50 nM Arp2/3, which is 100-fold lower than the VASP concentration of 5 μM. These results further demonstrate that the inclusion of Arp2/3 and VCA greatly reduces the ability of actin polymerization to deform VASP droplets (Figure 2g).

### Decreased droplet deformation is specifically due to Arp2/3 activity

We next investigated the molecular mechanisms by which Arp2/3 and VCA inhibited actin bundling. The first possibility we considered was that a reduction in actin partitioning to VASP droplets in the presence of Arp2/3 and VCA resulted in less actin polymerization within droplets and therefore a reduction in high aspect ratio structures. A reduction in partitioning could occur if Arp2/3 and VCA were to compete with actin for space within the crowded droplet environment. To evaluate this idea, we used the data in Figures 2a,b to plot the droplet aspect ratio as a function of the actin intensities inside the droplets, Figure 3a. This plot demonstrates that aspect ratio declines substantially with increasing Arp2/3 concentration, even when droplets with the same actin intensity are compared. These results indicate that steric exclusion of actin from VASP droplets in the presence of Arp2/3 and VCA cannot explain the loss of high aspect ratio structures and likely does not occur to a substantial degree.

**Figure 3.**
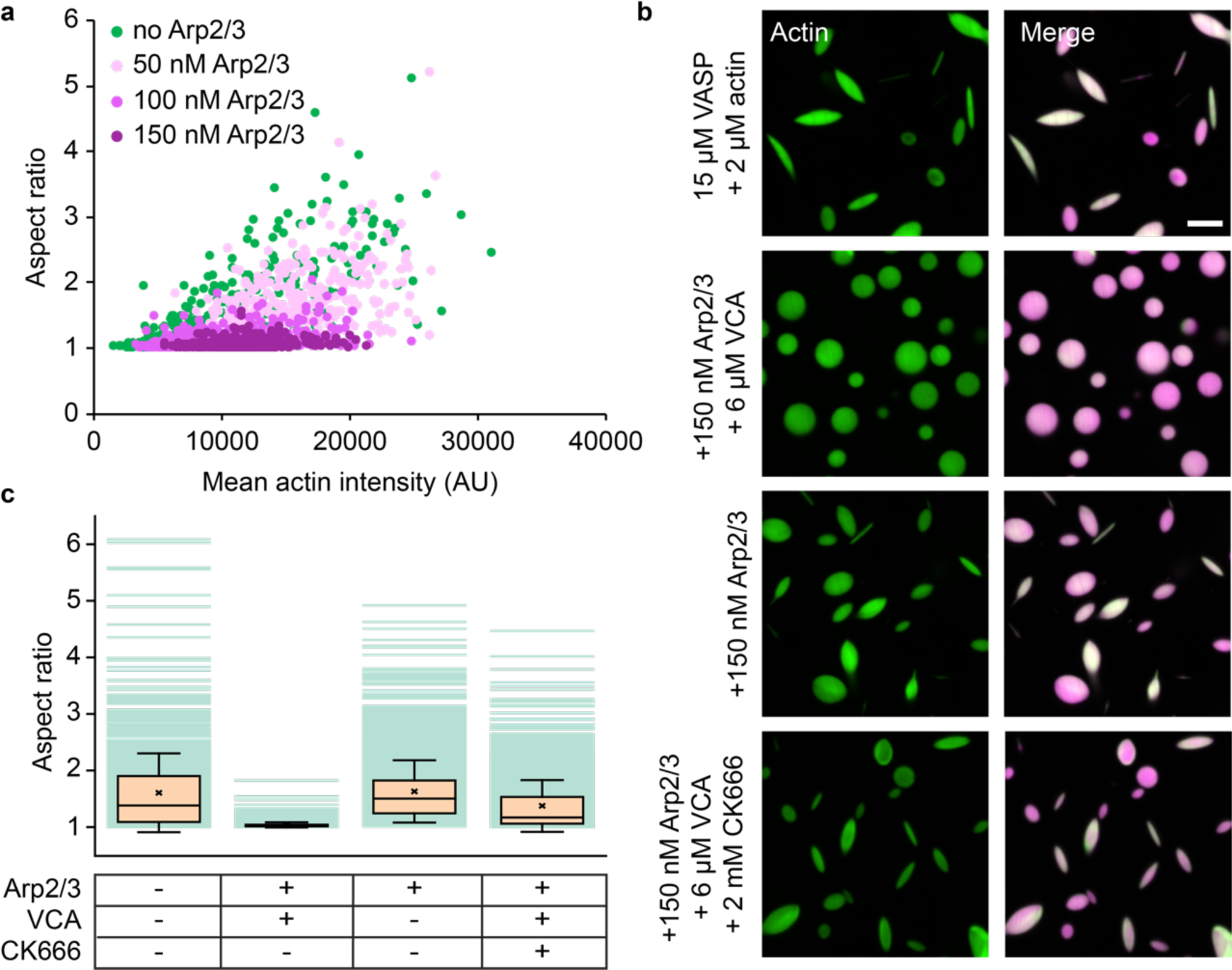
Decreased droplet deformation is due to Arp2/3 activity. (a) Relationship between droplet aspect ratio and actin intensity within the droplet for droplets formed from 15 μM VASP and exposed to 2 μM actin with increasing concentrations of Arp2/3 and VCA, as visualized in Figure 2a. Replicates can be found in Supplemental Figure S1. n=2885 droplets counted over at least 3 images per condition. (b) Representative images of droplets composed of 15 μM VASP and 2 μM actin, with either (i) Arp2/3 and VCA added, (ii) only Arp2/3, or (iii) Arp2/3, VCA, and the Arp2/3 inhibitor, CK666. Scale bar 5 μm. (c) Quantification of droplet aspect ratio measured from (b). Box represents IQR with median as a bisecting line, mean as an ‘x’, and whiskers represent SD. Data from 3 replicates. At least 700 droplets were analyzed per condition.

We next sought to determine whether the ability of Arp2/3 and VCA to nucleate assembly of branched actin networks could explain the reduction in droplet aspect ratio. Since Arp2/3 is activated by the WASP VCA domain, we tested the impact of omitting VCA on droplet aspect ratio, Figure 3b. The resulting droplets, which contained inactive Arp2/3, had a comparable distribution of aspect ratios to those consisting only of VASP and actin, demonstrating that activation of Arp2/3 is required for the decrease in aspect ratio (Figure 3b,c). To further elucidate the role of actin branching, we assayed droplets of VASP, Arp2/3, VCA, and actin in the presence of the Arp2/3 inhibitor, CK666 (31,32). Similar to the absence of VCA, inhibition of Arp2/3 by CK666 resulted in a distribution of aspect ratios similar to those of droplets consisting only of VASP and actin (Figure 3b,c). Together, these data show that activation of Arp2/3, which is known to drive assembly of branched actin networks (30,33) is responsible for the decrease in aspect ratio of droplets.

### Arp2/3 activity reduces droplet deformation by inhibiting assembly of actin rings

We next sought to determine the mechanism by which Arp2/3 activity inhibits droplet deformation. For this purpose, we examined the architecture of actin filaments within droplets. First, we repeated the experiment in Figure 2a, but stained with phalloidin, which specifically binds to actin filaments, rather than directly labeling actin monomers (Figure 4a). Our previous work showed that when actin polymerizes in droplets consisting only of VASP, actin filaments frequently partition to the inner periphery of droplets, where their curvature is minimized (10). In contrast, we found that, as the Arp2/3 and VCA fraction increased, actin filaments were less likely to adopt peripheral actin distributions, instead being distributed throughout the droplet (Figure 4a,b). Specifically, we found that droplets with an increased fraction of Arp2/3 and VCA showed less accumulation of actin at the inner surfaces of droplets (Figure 4b). Further, Arp2/3 and VCA containing droplets that displayed peripheral actin accumulation tended to have a more diffuse, “fuzzy” appearance (Figure 4c). More quantitatively, as the concentration of Ap2/3 increased, actin partitioning to the periphery was reduced, leaving a higher concentration of actin in the interior of the droplet (Figure 4c,d).

**Figure 4.**
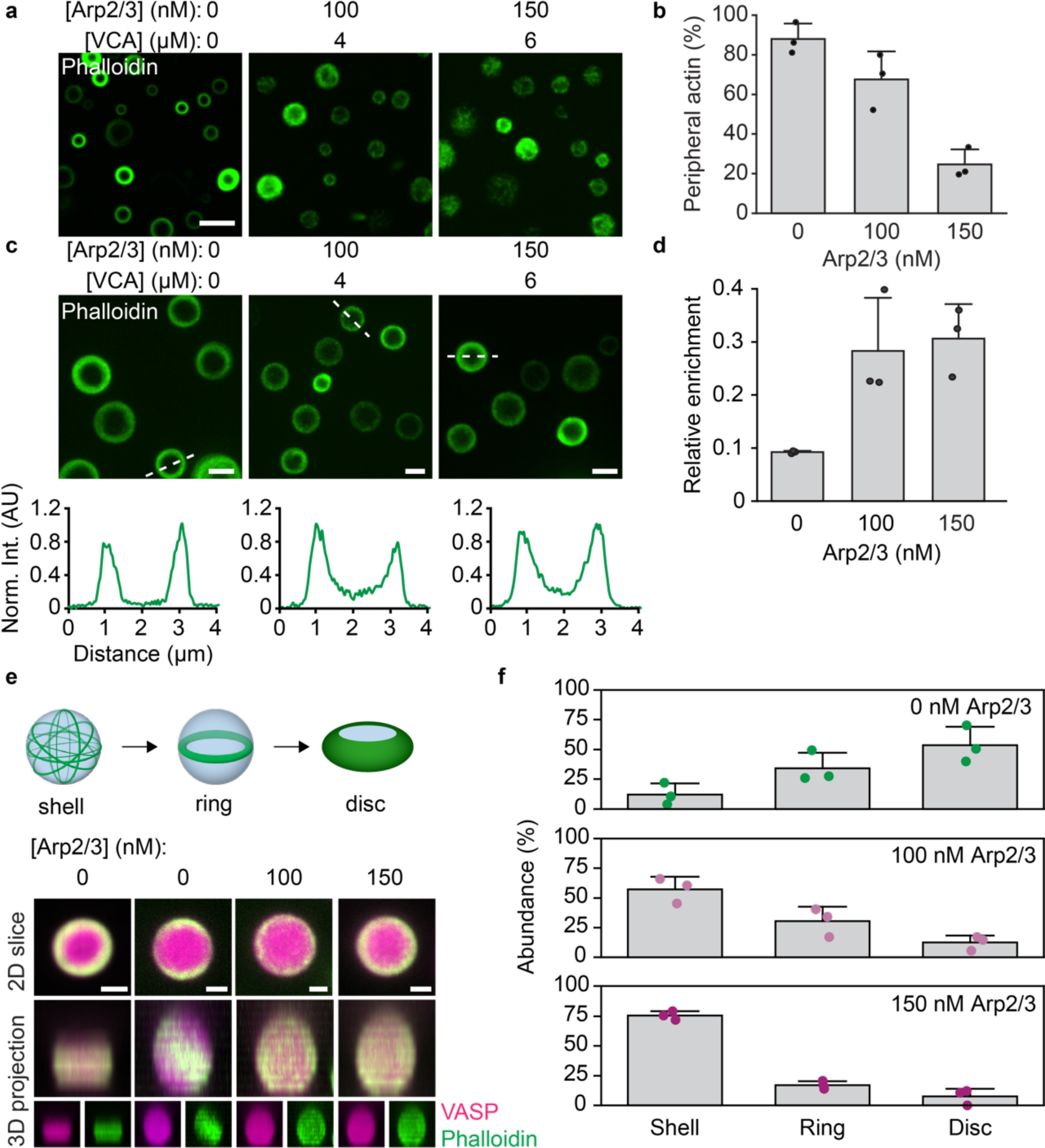
Arp2/3 activity reduces droplet deformation by inhibiting assembly of actin rings. (a) Representative images of droplets formed from 15 μM VASP with 2 μM actin and increasing concentration of Arp2/3 and VCA, stained with phalloidin. Scale bar 5 μm. (b) Quantification of the abundance of droplets with peripheral actin accumulation as a function of Arp2/3 concentration. Bars represent mean±SD of n=3 experiments, with 3 images analyzed per condition. (c) *Top:* Representative images of droplets formed as in (a), exhibiting peripheral accumulation of actin filaments. Scale bar 2 μm. *Bottom:* Representative line profiles of actin intensity in designated droplets. Actin intensities are normalized to the max intensity of each profile. (d) Quantification of the relative enrichment of actin to the interior of the droplet compared to the periphery. Data are mean±SD from n= 3 experiments, with 12 droplets analyzed per condition. (e) *Top:* Cartoon of the different 3D spatial arrangements of droplets with peripheral actin. *Bottom:* Representative 3D projections of droplets as in (c), revealing examples of disc, ring, and shell distributions of actin. Scale bar 1 μm. (f) Distribution of the abundance of actin shells, rings, and discs in droplets formed from 15 μM VASP with 2 μM actin and increasing Arp2/3 and VCA concentrations as shown in (a-d). Bars are mean±SD across n=3 experiments, with 3 images analyzed per condition.

In droplets consisting of VASP alone, once actin filaments accumulate at the droplet periphery, they spontaneously rearrange into a planar ring that gradually overcomes the droplet surface tension, flattening spherical droplets into discs (Figure 4e, top) (10). We analyzed 3D reconstructions of the data in Figure 4a,c to determine the impact of Arp2/3 and VCA on these transitions. The reconstructed images show that as the concentration of Arp2/3 increased, droplets mainly contained peripheral actin “shells”, which largely failed to condense into two-dimensional rings and discs (Figure 4e,f). These data indicate that Arp2/3 activity inhibits the rearrangement of actin filaments into the higher order structures required for droplet deformation. Specifically, a decrease in assembly of actin rings would fundamentally undermine droplet deformation and actin bundling. Without the bundling of actin into more compact and rigid structures, the droplet is less likely to deform into an elongated shape.

### Simulations suggest that filament branching inhibits the formation of actin-rich rings by reducing the average length of filaments

Our experiments demonstrate that Arp2/3 activity is responsible for inhibiting assembly of actin-rich rings inside VASP droplets, but they do not resolve the changes in the morphology of the actin network that are responsible for this inhibition. To gain greater insight into these phenomena, we used Cytosim, an agent-based framework for modeling cytoskeletal dynamics (34), to perform mechanochemical simulations of actin networks within droplets with a fixed radius of 1 μm. To capture their mechanical deformability, actin filaments were discretized into a series of segments that were allowed to bend along hinge points (Supplemental figure S2). Actin, VASP, and Arp2/3 molecules were constrained within the boundary of spherical droplets using harmonic potentials (Supplemental figure S2). Additionally, spatial overlap between filaments and VASP molecules was prevented by applying steric potentials (Supplemental figure S2). Chemical evolution of the network in Cytosim involved actin filament growth, binding, and unbinding of VASP to actin, and binding of Arp2/3 to actin, resulting in nucleation of filament branches (Supplemental figure S2). The simulations also accounted for Brownian dynamics of the molecules considered. While kinetic parameters for Arp2/3 were chosen from experimental data (Supplementary Table S3), kinetic parameters for VASP were chosen based on simulations of actin polymerization within VASP droplets that lacked Arp2/3 (35). The present simulations are described in detail in the Supplementary Methods section, Supplementary Figure S2, and Supplementary Table S3.

Figure 5a shows snapshots of the simulation after 10 minutes, which provided sufficient time for the distribution of actin inside the droplets to approach steady state. In the absence of Arp2/3, actin rings form at the inner periphery of the droplets, in agreement with experimental results (Figure 4e). However, as the concentration of Arp2/3 increased, we observed an increasing number of actin filaments within the droplet that were not part of the ring. Specifically, at Arp2/3 concentrations above 100 nM, the actin distribution within the droplet became largely homogenous. To determine the radial distribution of F-actin within the droplet, we discretized the volume into a series of overlapping shells of finite thickness (25 nm). Computing the ratio of local F-actin density to the bulk F-actin density for each of these shells, we observed that the ring configuration in the absence of Arp2/3 displayed a marked accumulation of actin at the periphery of the droplet and was devoid of actin elsewhere (Figure 5b). As the concentration of Arp2/3 increased, we observed three distinct regions within the droplets, each of which had different actin distributions. Near the center of the droplet (r ≤ 0.1 μm) the density ratio was dominated by small volume effects and is therefore not depicted in Figure 5b. In the droplet core (0.1 < r ≤ 0.6), we observed a steady increase in the density ratio as the concentration of Arp2/3 increased, which is summarized in Figures 5b,c. Finally, in the droplet periphery (r > 0.6), we observed a marked decrease in actin accumulation. These distinct behaviors can be understood by looking at changes in the characteristics of the filaments with increasing Arp2/3 concentration. Specifically, Figure 5d shows that increasing the concentration of Arp2/3 resulted in enhanced nucleation of actin filaments throughout the trajectory. As a result, the median filament length decreased, resulting in an increasing number of filaments with lengths below the droplet diameter (Figure 5e). These shorter filaments do not have to bend to remain within the confines of the droplet, and as a result, their driving force to partition to the droplet periphery is greatly reduced (Figure 5f).

**Figure 5.**
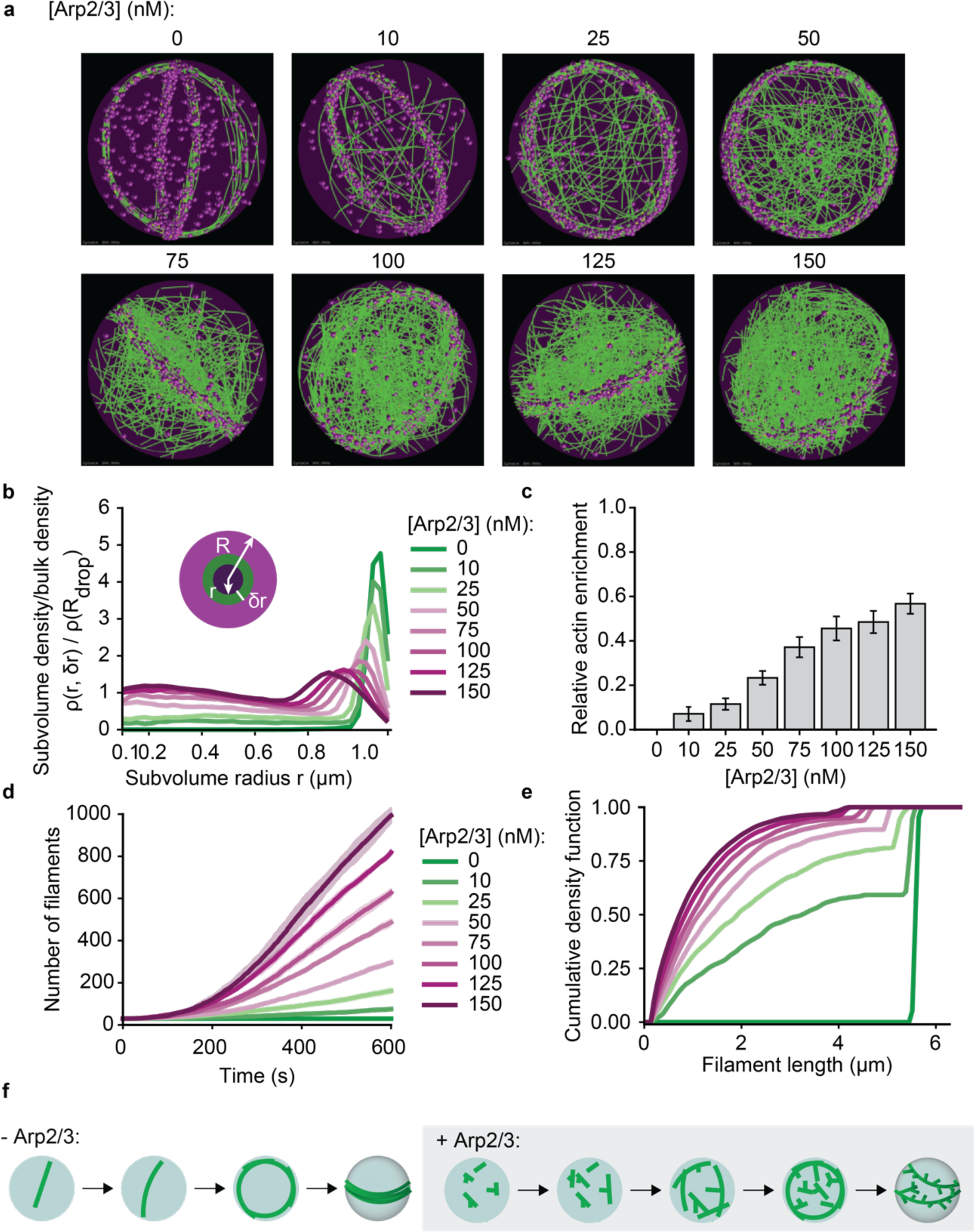
Simulations predict that Arp2/3-driven changes to filament length distribution inhibits the formation of actin-rich rings at the droplet periphery. (a) Representative final snapshots (t=600s) from rigid droplets (radius = 1 μm) [VASP-tet] = 0.2 μM and various Arp2/3 concentrations are shown. Actin filaments are shown in green while VASP tetramers are shown as magenta spheres. Supplemental Movie M3 shows the time evolution of the trajectories. (b) Plot shows the mean ratio of actin density within subvolume shells of radius r and thickness *δr* = 25 *nm* (shown in inset) to the overall density of actin within the droplet of radius R. Plot lines are colored based on Arp2/3 concentration (c) Actin enrichment close to droplet core is calculated at various [Arp2/3] as the ratio of actin density within shells of radius *r_c_* = 0.25 *μm* and *r*_#_ = 0.9 *μm*. Data from 5 replicates, the last 10 snapshots per replicate were used here. (d) Mean (solid line) and standard deviation (shaded area) in the number of actin filaments are plotted as time traces and colored based on the concentration of Arp2/3. (e) The final distribution of filament lengths (last 10 snapshots per replicate) is plotted as a cumulative density function and colored by Arp2/3 concentration. Data from 5 replicates. (f) Proposed model describing ring formation in the absence and presence of Arp2/3 activity.

Taken together, the results of our experiments (Figure 4) and simulations (Figure 5) suggest that the ability of filament branching to undermine ring formation can be explained by two key phenomena. First, branches increase the effective diameter of filaments, sterically inhibiting filament bundling within rings. This finding is consistent with previous studies of actin-Arp2/3 interactions (36,37). Second, branched actin filaments are shorter, on average, than unbranched filaments, because the actin monomers within the branch are unavailable to add to the length of the mother filament. The resulting shorter filaments cannot exert sufficient forces to overcome the surface tension of the droplet. Specifically, when the length of filaments exceeds the droplet diameter, filaments must bend to remain within the droplet. Because the persistence length of actin, ∼17 μm (38), is greater than the diameter of most of the droplets in our experiments (10) (Figure 1c), this bending comes at a substantial energetic cost that can be minimized by partitioning filaments to the inner droplet surface, where their curvature is lowest (39). Since branched filaments are fundamentally shorter, they will be less likely to encounter this constraint, when compared to longer, linear filaments.

### At high actin-to-VASP ratios, Arp2/3 activity drives aster-like morphologies

Up to this point, our data indicate that for relatively low actin to VASP ratios, Arp2/3 activity strongly inhibits droplet-mediated actin bundling. In particular, results from experiments, further elucidated by simulations, suggest that Arp2/3 activity results in an abundance of short, branched filaments that fail to deform droplets. Similarly, prior work has shown that branching and bundling proteins compete with one another when the supply of actin is limited (40). Together, these results suggest that a limited supply of actin may be inhibiting droplet deformation in our experiments. Based on this reasoning, we next explored the impact of increasing the actin concentration in our system. Specifically, we added increasing concentrations of Arp2/3 to droplets formed at higher actin to VASP ratios, beginning with a 1:1 ratio (5 μM VASP and 5 μM actin), holding VCA concentration constant at 4 μM (Figure 6a). At Arp2/3 concentrations as low as 50 nM, the VASP droplets began to deform into forked structures, which we did not observe at the lower actin to VASP ratios used in Figure 2. These forked structures increased in abundance as the Arp2/3 concentration increased.

**Figure 6.**
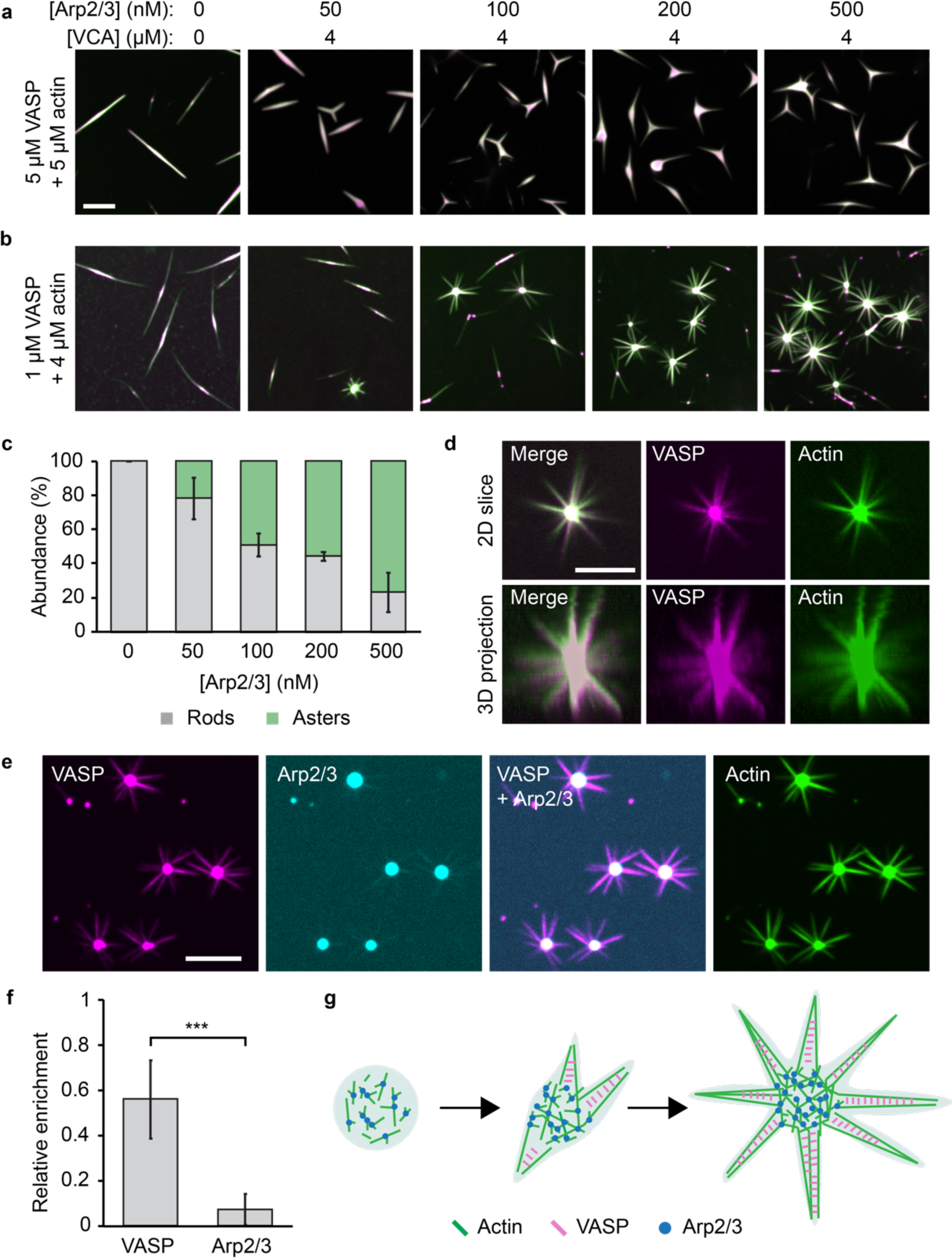
At high actin-to-VASP ratios, Arp2/3 activity drives aster-like morphologies. (a) Titration of Arp2/3 and 4 μM VCA in droplets formed from 5 μM VASP and exposed to 5 μM actin. Scale bar 5 μm. (b) Titration of Arp2/3 and 4 μM VCA as in (a) into droplets formed from 1 μM VASP and exposed to 4 μM actin. Scale bar 5 μm. (c) Quantification of the percent of droplets that deformed into rods and asters in (b). Bars are mean±SD across n=3 experiments, with at least 4 images analyzed per condition. (d) 3D projection of a representative aster-shaped droplet. Scale bar 5 μm. (e) Representative images of VASP, Arp2/3, and actin intensity distributions throughout aster-shaped droplets. Condition: 1 μM VASP, 500 μM Arp2/3, 4 μM VCA, 4 μM actin. Scale bar 5 μm. (f) Quantification of the relative enrichment of VASP and Arp2/3 to the spokes of aster-shaped droplets. Enrichment is normalized to actin intensity. Droplets formed as in (e). Bars are mean±SD across n=3 experiments, with 10 droplets analyzed per experiment. Asterisks indicates data tested for significance via an unpaired two tailed t-test, with p < 0.001. (g) Proposed model for Arp2/3- and VASP-mediated aster formation.

Next, we further increased the actin to VASP ratio to 4:1 (1 μM VASP and 4 μM actin), again, gradually increasing the concentration of Arp2/3, while holding VCA concentration constant at 4 μM (Figure 6b). At this actin to VASP ratio, multi-pronged, three-dimensional, aster-like structures emerged with increasing Arp2/3 concentration (Figure 6c,d). These droplet-mediated asters appear similar to previously reported actin asters, which formed on beads coated with WASP, Arp2/3, and fascin, an actin bundling protein (41). In these previous studies, Arp2/3 at the surface of the bead nucleated a branched actin network, which was then bundled by fascin to form radiating spokes. Similar results have also been observed in the absence of beads, in which actin self-assembled into asters in the presence of VCA, Arp2/3, and fascin (29,42,43). Importantly, our system lacks fascin. However, because VASP is a tetramer, it is able to crosslink actin filaments, such that it acts as a bundling protein (16,44).

Interestingly, in droplet-mediated asters, we found that VASP colocalized with actin in both the droplet core and along the spokes, while Arp2/3 was mainly localized to the droplet core and depleted from the spokes (Figure 6e,f). Specifically, comparison of the spoke to core intensity ratio for VASP and Arp2/3 showed that VASP’s relative partitioning to the spokes was nearly 10-fold greater than Arp2/3’s (partitioning of 6.8±2.3% compared to 0.69±0.52%) (Figure 6f). This result suggests that branched actin structures are more likely to exist in the core of the asters, while the spokes likely consist of more linear filaments, bundled by VASP, Figure 6g. In this way, the protein droplet, which consists of VASP, Arp2/3 and VCA, mediates the segregation of branched and bundled actin networks.

## DISCUSSION

We have previously shown that liquid-like protein droplets consisting of VASP can catalyze the polymerization of actin and its assembly into parallel bundles (10). In the present work, we show that creation of a branched actin network, through the activity of Arp2/3, competes with the filament bundling activity of VASP. In particular, at low actin to VASP ratios, introduction of Arp2/3 resulted in potent inhibition of droplet bundling, such that droplets remained spherical (Figure 2). The resulting branched network of filaments failed to break symmetry, forming a relatively homogenous actin mesh within the droplets, rather than accumulating at the droplet periphery (Figure 4,5). Computational modeling suggested that this lack of filament partitioning to the droplet periphery is due to a reduction in filament length, as well as an increase in steric hindrance among branched filaments. Guided by these results, we found that at higher actin to VASP ratios, bundles emerged from the droplets, resulting in aster-like morphologies.

Interestingly, recent work showed that, when Arp2/3 and actin are co-encapsulated in vesicles, actin branching suppresses fascin-mediated bundling (40). This observation was attributed to competition between fascin and Arp2/3 for a limited supply of actin. This competition may parallel the competition that we have observed between VASP and Arp2/3. Specifically, when the concentration of actin in our droplet system increased, bundling re-emerged, resulting in forked and aster-like structures. These structures likely formed because the increased actin concentration allowed the growth of longer actin filaments, which are more easily bundled. Interestingly, VASP was enriched in the spokes of these structures, while Arp2/3 was enriched in their cores, further suggesting that the spokes consisted primarily of bundled actin, which emerged from a branched actin core. This observation parallels prior studies that have reported the spontaneous formation of actin asters in the presence of Arp2/3, WASP, and fascin (29,41,42). In all systems, a branched actin core, rich in Arp2/3, gave rise to bundled filaments that were rich in actin bundling proteins– either fascin or VASP.

How does an actin bundle emerge from a network of branched filaments? One hypothesis is that actin filaments, nucleated by Arp2/3, need to reach a critical length before they can become bundled (29,45). Specifically, filaments need to be long enough to bend toward each other so that they can be crosslinked by a bundling protein, such as VASP (29). Longer filaments would likely concentrate at the droplet periphery, where their curvature is minimized. Assembly of these peripheral filaments into bundles could explain the aster-like morphologies that we observe (Figure 6). Additionally, the droplet’s surface tension may enhance bundling by opposing filament branching, similar to previous observations in reconstituted membrane systems (46). In this way, the mechanics of the droplet could promote the emergence of actin bundles that originate from a branched actin interior.

At the leading edge of the cell, the sheet-like lamellipodium is filled with a branched actin network that relies on Arp2/3-mediated nucleation (47). Filopodia, finger-like membrane protrusions rich in bundled actin, emerge from the lamellipodium through the action of actin polymerases, as well as bundling proteins. Growth of filopodia requires that long, unbranched filaments emerge from the branched lamellipodium (4,8). VASP has been shown to assist in this process by i) processively elongating filaments, ii) clustering barbed ends, and iii) bundling filaments (44,48,49). Importantly, VASP localizes to the tips of filopodia (12,50,51). In this way, filopodia are enriched in VASP and depleted of Arp2/3. Interestingly, this finding parallels the segregation of VASP and Arp2/3 in our droplet-mediated asters, suggesting that protein condensation could play a role in mediating branch-to-bundle transitions of the cytoskeleton (Figure 6g).

Overall, this work suggests new roles for phase separation in the organization of the cytoskeleton. The cellular leading edge is inherently dynamic, owing to the continual rearrangement of the membrane, actin, and associated machinery. Importantly, Arp2/3 and WASP-family proteins have been found to participate in protein condensation (26–28), and oligomerization of WASP family proteins synergistically increases their activity (52,53). VASP has also been shown to form dynamic clusters on the lamellipodial membrane that exhibit fusion and fission (14), and clustering of VASP enhances its processivity as an actin polymerase (48). While the role of protein condensation in the cytoskeleton is still emerging, these results collectively suggest multiple mechanisms by which condensates could participate in network remodeling. In particular, our work suggests that, by tuning the composition of condensates, cytoskeletal networks can dictate the assembly of branched or bundled architectures.

## Acknowledgements

This research was supported by grants from the National Institutes of Health to JCS (R35GM139531) and PR (R01GM132106), by the National Science Foundation through a Modulus Grant BIO-1934411 to PR and JCS, and by the Welch Foundation through Grant F-2047 to JCS. AC would like to acknowledge Dr. Francois Nedelec for feedback on the model schematic.

## Author contributions

KG, AC, EML, PR, and JCS designed experiments. KG, AC, PR, and JCS wrote and edited the manuscript. KG, NY, AC, and LW performed experiments and analyzed data. All authors consulted on manuscript preparation and editing.

## Competing interests statement

The authors declare no competing interests.

## METHODS

### Reagents

Porcine brain Arp2/3 complex, human GST-tagged N-WASP VCA domain, rabbit skeletal muscle actin, and ATP were purchased from Cytoskeleton. Tris base, CaCl2, TCEP, NaCl, KCl, MgCl2, and poly-L-lysine, and PEG 8000 were purchased from Sigma-Aldrich. Alexa Fluor 647 C2 maleimide was purchased from Thermo Fisher Scientific. Maleimide-Atto 488 and maleimide-Atto 594 were purchased from ATTO-TEC. Phalloidin-iFluor-488 and Arp2/3 inhibitor CK666 were purchased from Abcam. mPEG-SVA (MW 5000) was purchased from Laysan Bio.

A “cysteine-light” mutant form of human VASP was expressed from a pET vector (pET-6xHis-TEV-KCK-VASP(CCC-SSA)) that was a gift from Scott Hansen (23). Expression, purification, and cleavage of the His-tag from the VASP protein was conducted as explained previously (10). The resulting purified protein was stored in 25 mM HEPES pH 7.5, 200 mM NaCl, 5% Glycerol, 1 mM EDTA, 5 mM DTT. Single-use aliquots were flash frozen and stored at −80°C until the day of an experiment.

### Protein labeling

The “cysteine-light” form of VASP allows for selective labeling of the single N-terminal cysteine using maleimide-conjugated dyes, as performed previously (10). Briefly, VASP was incubated with a 3-fold molar excess of dye for 2 hours at room temperature to achieve a 10-20% labeling ratio. The protein was separated from the free dye by applying the labeling reaction to a Princeton CentriSpin-20 size exclusion column (Princeton Separations). Labeled VASP was eluted and stored in 50 mM Tris pH 7.8, 300 mM NaCl, 0.5 mM EDTA, 0.5 mM EGTA, 5 mM TCEP. Monomeric actin was labeled on Cys-374 using maleimide-Atto 488. Dye was incubated with G-actin at a 3-5 fold molar excess for 2 hours at room temperature to achieve a labeling ratio of 20-30%. Unconjugated dye was separated from labeled actin by applying the labeling reaction to a Princeton CentriSpin-20 size exclusion column hydrated with A buffer (5 mM Tris-HCl pH 8, 0.2 mM CaCl_2_, 0.2 mM ATP, 0.5 mM DTT). The eluted labeled protein was then centrifuged at 14k rpm for 15 min at 4°C to remove aggregates, and flash-frozen in single-use aliquots. VCA and Arp2/3 were labeled with maleimide Atto-488 and maleimide Atto-594 respectively, following the same protocol for VASP but with the following minor modifications (5,17). Labeled VCA was eluted into 20 mM Tris pH 7.5, 25 mM KCl, 1 mM MgCl_2_, 0.5 mM EDTA, 2% sucrose. Arp2/3 was eluted into 10 mM imidazole pH 7, 50 mM KCl, 1 mM MgCl_2_, 1 mM EGTA, 0.5 mM DTT, 100 μM ATP, 5% sucrose. After labeling, both proteins were centrifuged at 21k rpm for 15 min at 4°C to remove aggregates, and used the same day.

### VASP droplet formation and actin polymerization

VASP droplets were formed as described previously (10). Briefly, VASP was mixed with “droplet buffer”, 3% (w/v) PEG(8k) in 50 mM Tris pH 7.4, 150 mM NaCl, 5 mM TCEP to the final concentrations stated in each experiment. PEG was added last to induce droplet formation. For experiments with Arp2/3 and VCA, these proteins were mixed with VASP in droplet buffer at the desired concentrations, then PEG was added to induce droplet formation.

For droplet polymerization experiments with labeled monomeric actin, actin was added after the droplets were allowed to form for 10 minutes. Actin was allowed to polymerize in the droplets for 15 minutes, then the droplets were imaged. For experiments with phalloidin-stained actin, unlabeled monomeric actin was added to preformed VASP droplets, allowed to polymerize for 15 minutes, then phalloidin was added to stain the filaments for another 15 minutes before imaging.

### Microscopy

Samples were prepared in wells formed by 1.5 mm thick silicone gaskets (Grace Biolabs) on Hellmanex II (Hellma) cleaned, no.1.5 glass coverslips (VWR) passivated with poly-L-Lysine conjugated PEG5k. A second coverslip was placed on top to seal the imaging chamber to prevent evaporation during imaging. Fluorescence microscopy was performed using the Olympus SpinSR10 spinning disc confocal microscope fitted with a Hamamatsu Orca Flash 4.0V3 SCMOS Digital Camera. FRAP was performed using the Olympus FRAP unit 405 nm laser.

PLL-PEG was prepared as described previously with minor alteration (54). Briefly, amine-reactive mPEG-SVA was conjugated to Poly-L-Lysine (15-30 kD) at a molar ratio of 1:5 PEG to poly-L-lysine. The conjugation reaction was performed in 50 mM sodium tetraborate pH 8.5 solution and allowed to react overnight at room temperature with continued stirring. The product was buffer exchanged into PBS pH 7.4 using Zeba spin desalting columns (7K MWCO, ThermoFisher) and stored at 4°C.

### Image analysis

ImageJ was used to quantify the distribution of droplet characteristics. Specifically, droplets were selected by thresholding in the VASP channel, and shape descriptors (e.g. diameter, aspect ratio) and protein intensities were measured using the analyze particles function.

FRAP data were analyzed using the FRAP Profiler plugin for ImageJ. Fluorescence recovery of a 2.5 μm diameter region was measured over time, and intensities were normalized to the maximum pre-bleach and minimum post-bleach intensity. All intensity data was corrected for photobleaching. Recovery was measured only for droplets of similar diameters.

Quantification of peripheral actin abundance was performed by scoring phalloidin-stained droplets for the presence of peripheral actin accumulation. Specifically, droplets were considered to have peripheral accumulation of actin if there was a decrease in actin fluorescence intensity in the center of the droplet, as verified by line profiles drawn across the droplet diameter. Droplets that had fused to other droplets were omitted. Droplets containing peripheral accumulation of filamentous actin were further classified based on their three-dimensional arrangement of actin: either shells, rings, or discs, as performed previously (10). Rings, discs, and shells were differentiated by making 3D projections of each individual droplet. Shells were defined as spherical VASP droplets that have uniform spherical actin intensity in the 3D projection. Rings were defined as spherical VASP droplets that contained a planar arrangement of actin in the 3D projection. Discs were defined as non-spherical, planar droplets that colocalized both actin and VASP. Only droplets with peripheral accumulation of actin in the imaging plane were counted.

To calculate the partitioning of actin to the center of the droplet compared to the droplet periphery, the average intensity of a 15×15 pixel square in the center of the droplet was measured. This value was divided by the average intensity of the actin-rich ring formed at the droplet periphery to derive the relative enrichment in the center. The average intensity of the ring was measured by performing a radial transformation originating at the center of each droplet, and then measuring the average actin intensity along a line profile drawn in the center of each actin ring. All droplets analyzed were randomly selected.

To quantify the abundance of asters, images were scored for either rod-shaped droplets or aster-shaped droplets. Aster-droplets were defined as droplets with at least 4 actin-rich spokes radiating off of the central droplet core. Rods were defined as 2D linear droplets with VASP colocalized along the entire length. Droplets that intersected or overlapped with other droplets were not counted.

To calculate the partitioning of VASP and Arp2/3 to the aster-droplet spokes, the average intensities of the proteins in the spokes were divided by the average intensities in the droplet core, then normalized to the actin channel. Specifically, in a single z-slice in which the droplet core and spokes were in focus, the average intensities of Arp2/3, actin, and VASP in the droplet core were measured using ImageJ as mentioned above. To measure the intensity of proteins in the spokes, a circular line profile 1 μm from the surface of the droplet core was drawn, perpendicular to the spokes. Each intensity peak in the line profile corresponded to a spoke radiating off of the droplet. Using the actin channel as the reference for spoke peaks, the corresponding spoke intensities in the VASP and Arp2/3 channel were measured, and the resulting average spoke intensity was used to calculate the average spoke partitioning per droplet. Importantly, labeled VASP and Arp2/3 were used separately to avoid spectral overlap and bleed through.

## SUPPLEMENTAL INFORMATION

**Supplemental Figure S1.**
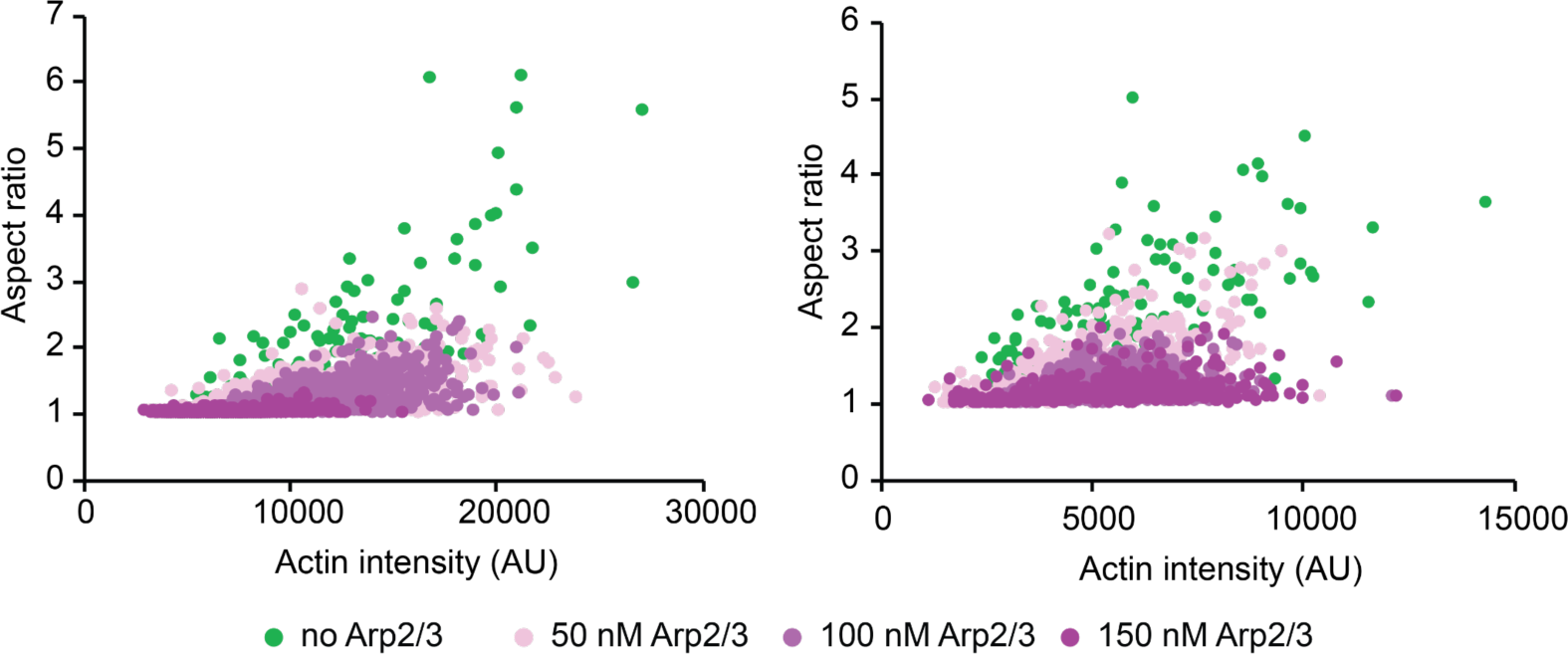
Replicates of figure 3a. Relationship between droplet aspect ratio and actin intensity within the droplet for droplets formed from 15 μM VASP and exposed to 2 μM actin with increasing concentrations of Arp2/3 and VCA, as visualized in Figure 2a. *Left:* n = 3387 droplets. *Right:* n= 2761 droplets.

**Supplemental Figure S2.**
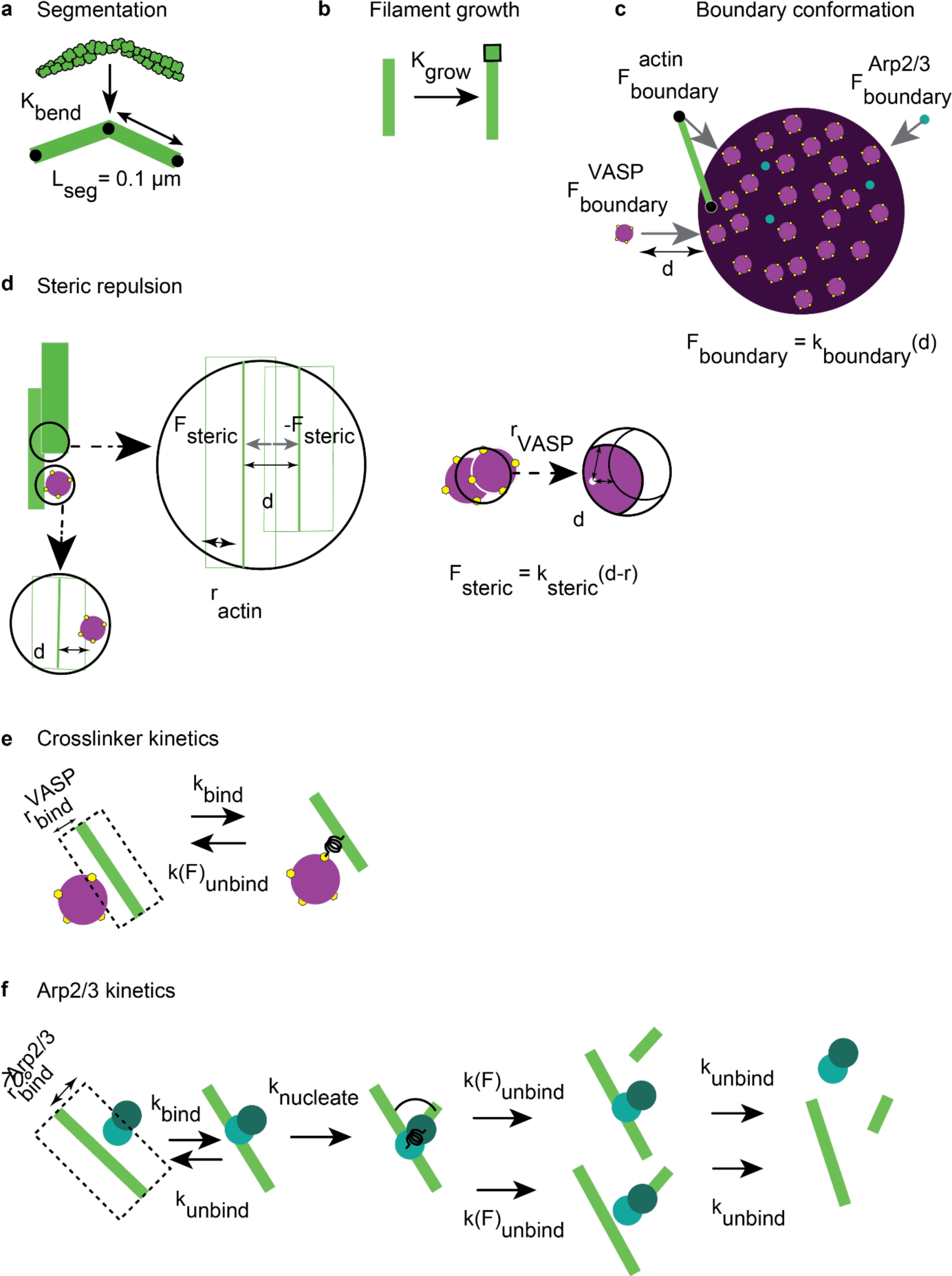
Depictions of model parameters. (a) Actin filament curvature is approximated through a series of line segments of length L_seg_. (b) Propensity of filament growth is calculated from the product of the G-actin monomers available in the volume and the filament growth rate K_grow_. (c) Actin (green), VASP tetramers (purple spheres), and Arp2/3 (teal) molecules are constrained within the droplet volume through harmonic potentials. (d) Steric repulsion potentials prevent spatial overlap of molecules within the droplet. (e) VASP binding sites (shown as yellow hexagons) located within the binding distance r_bind_^VASP^ stochastically bind actin filaments at rate k_bind_. Force sensitive unbinding reaction is also included. (f) Diffusing Arp2/3 molecules within binding distance r_bind_^Arp2/3^ bind actin stochastically at rate k_bind_. Branched nucleation at rate k_nucleate_ results in an offspring filament at 70° angle with the parent filament. Force-sensitive unbinding reactions at rate kunbind are also considered. Please refer to Supplementary Table S3 for a detailed table of various parameters used in the model.

**Table S3.**
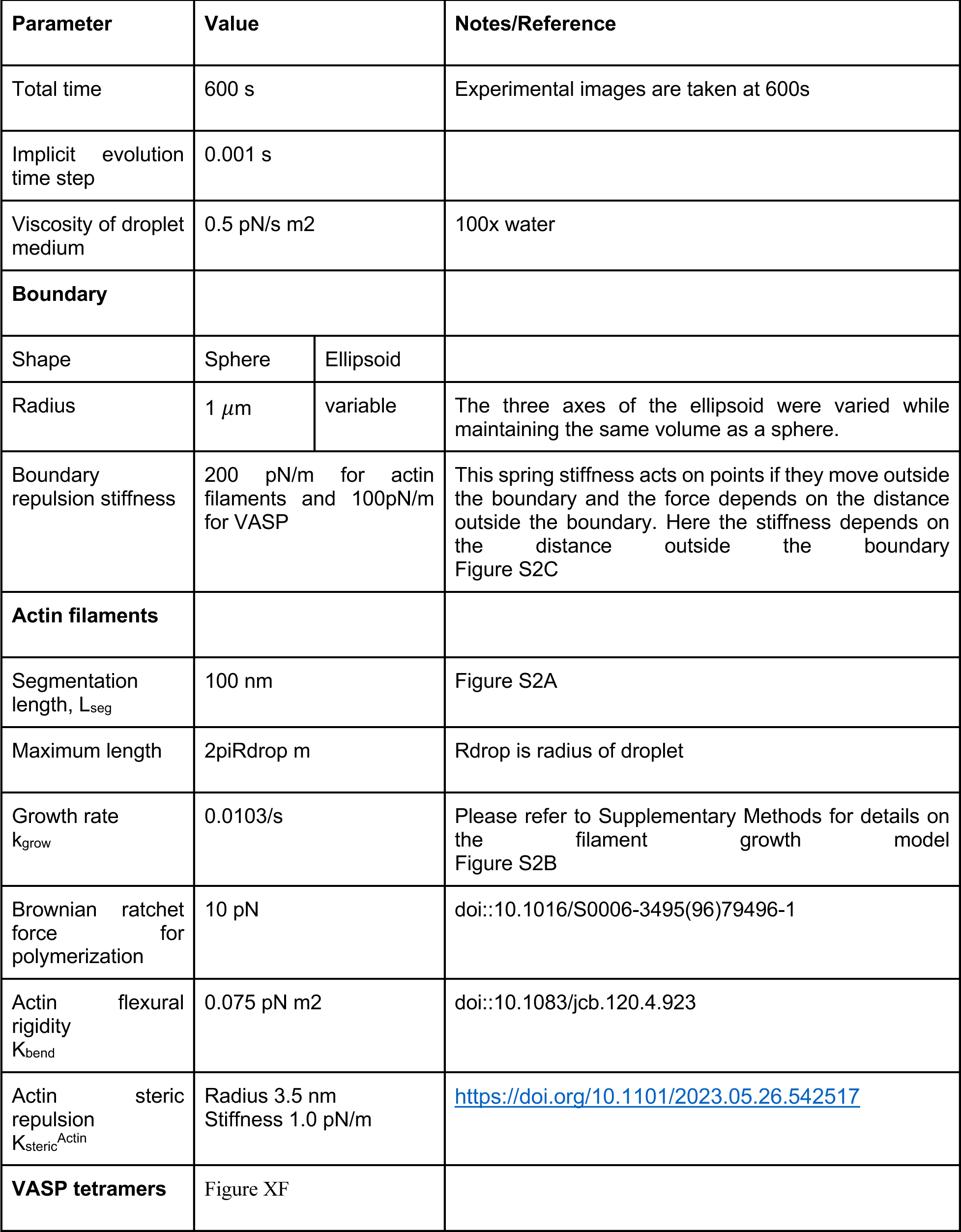
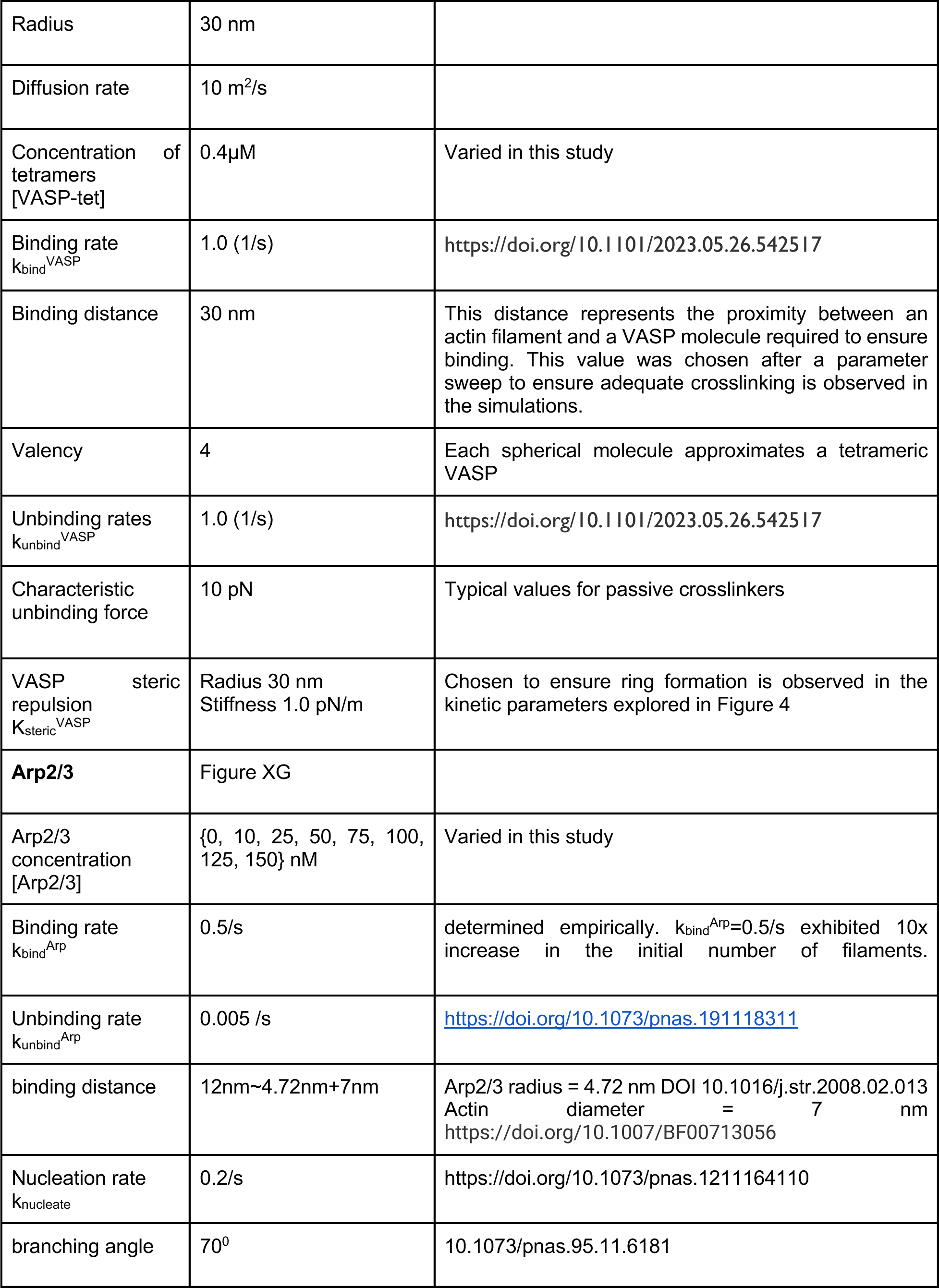
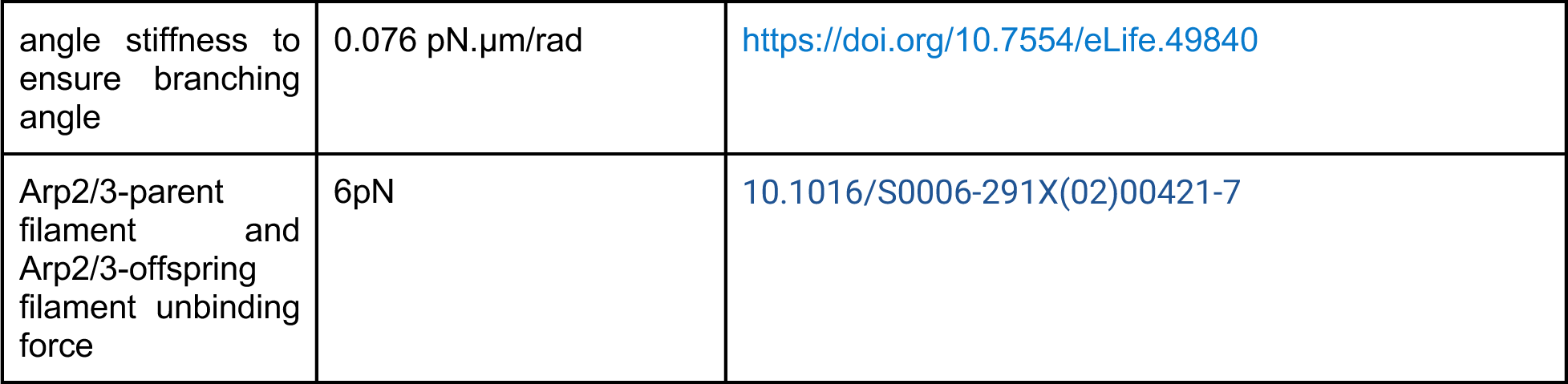
Table of parameters used to set up the actin model in Cytosim.

| Parameter | Value |  | Notes/Reference |
| --- | --- | --- | --- |
| Total time | 600 s |  | Experimental images are taken at 600s |
| Implicit evolution time step | 0.001 s |  |  |
| Viscosity of droplet medium | 0.5 pN/s m<sup>2</sup> |  | 100x water |
| **Boundary** |  |  |  |
| Shape | Sphere | Ellipsoid |  |
| Radius | 1 μm | variable | The three axes of the ellipsoid were varied while maintaining the same volume as a sphere. |
| Boundary repulsion stiffness | 200 pN/m for actin filaments and 100pN/m for VASP |  | This spring stiffness acts on points if they move outside the boundary and the force depends on the distance outside the boundary. Here the stiffness depends on the distance outside the boundary Figure S2C |
| **Actin filaments** |  |  |  |
| Segmentation length, $L_{seg}$ | 100 nm | | Figure S2A |
| Maximum length | $2\pi R_{drop}$ m | | $R_{drop}$ is radius of droplet |
| Growth rate $k_{grow}$ | 0.0103/s | | Please refer to Supplementary Methods for details on the filament growth model Figure S2B |
| Brownian ratchet force for polymerization | 10 pN |  | doi::10.1016/S0006-3495(96)79496-1 |
| Actin flexural rigidity $K_{bend}$ | 0.075 pN m<sup>2</sup> | | doi::10.1083/jcb.120.4.923 |
| Actin steric repulsion $K_{steric}^{Actin}$ | Radius 3.5 nm<br>Stiffness 1.0 pN/m | | https://doi.org/10.1101/2023.05.26.542517 |
| **VASP tetramers** | Figure XF |  |  |

|  |  |  |
| --- | --- | --- |
| Radius | 30 nm |  |
| Diffusion rate | 10 m<sup>2</sup>/s |  |
| Concentration of tetramers [VASP-tet] | 0.4μM | Varied in this study |
| Binding rate $k_{bind}^{VASP}$ | 1.0 (1/s) | https://doi.org/10.1101/2023.05.26.542517 |
| Binding distance | 30 nm | This distance represents the proximity between an actin filament and a VASP molecule required to ensure binding. This value was chosen after a parameter sweep to ensure adequate crosslinking is observed in the simulations. |
| Valency | 4 | Each spherical molecule approximates a tetrameric VASP |
| Unbinding rates $k_{unbind}^{VASP}$ | 1.0 (1/s) | https://doi.org/10.1101/2023.05.26.542517 |
| Characteristic unbinding force | 10 pN | Typical values for passive crosslinkers |
| VASP steric repulsion $K_{steric}^{VASP}$ | Radius 30 nm<br>Stiffness 1.0 pN/m | Chosen to ensure ring formation is observed in the kinetic parameters explored in Figure 4 |
| **Arp2/3** | Figure XG |  |
| Arp2/3 concentration [Arp2/3] | {0, 10, 25, 50, 75, 100, 125, 150} nM | Varied in this study |
| Binding rate $k_{bind}^{Arp}$ | 0.5/s | determined empirically. $k_{bind}^{Arp}=0.5/s$ exhibited 10x increase in the initial number of filaments. |
| Unbinding rate $k_{unbind}^{Arp}$ | 0.005 /s | https://doi.org/10.1073/pnas.191118311 |
| binding distance | 12nm~4.72nm+7nm | Arp2/3 radius = 4.72 nm DOI 10.1016/j.str.2008.02.013<br>Actin diameter = 7 nm<br>https://doi.org/10.1007/BF00713056 |
| Nucleation rate $k_{nucleate}$ | 0.2/s | https://doi.org/10.1073/pnas.1211164110 |
| branching angle | 70° | 10.1073/pnas.95.11.6181 |
| angle stiffness to ensure branching angle | 0.076 pN.μm/rad | https://doi.org/10.7554/eLife.49840 |
| Arp2/3-parent filament and Arp2/3-offspring filament unbinding force | 6pN | [10.1016/S0006-291X(02)00421-7](https://doi.org/10.1016/S0006-291X%2802%2900421-7) |

## SUPPLEMENTAL MOVIES

**Supplemental Movie M1.** VASP droplets formed with Arp2/3 and VCA still undergo fusion. **Supplemental Movie M2.** VASP droplets formed with Arp2/3 and VCA still exhibit fast and complete recovery after photobleaching.

**Supplemental Movie M3.** Simulation time-lapse of droplets with increasing Arp2/3 results in less bundled rings and more central actin accumulation.

## SUPPLEMENTAL METHODS

### Chemical and mechanical framework employed in CytoSim

Cytosim (https://gitlab.com/f-nedelec/cytosim) is an agent-based framework to simulate cytoskeletal networks with physical realism subject to geometric constraints. In Cytosim, each time step (1ms in this study) involves the evolution of chemical reactions based on their propensities, followed by Brownian dynamics evolution of the chemical species to capture diffusion effects. Filament extension is modeled as follows: Assuming a growth rate of 0.0103/s, we solved the following ordinary differential equations in series to identify the total actin parameter to use in Cytosim [T-Actin].

| Symbol | Represents concentration of |
| --- | --- |
| [P] | Plus End |
| [G-actin] | unpolymerized actin |
| [F-actin] | Polymerized actin |
| $N_{fil}$ | Number of filaments |

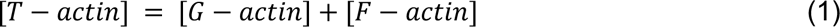

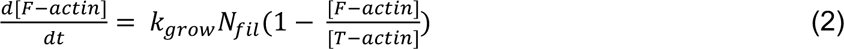

These equations were solved with initial conditions *t* = 0, *N_fil_* = 30, [*F* − *actin*](0) = 0.44 *μM* along with boundary conditions *t* = 600*s*, [*F* − *actin*](600) = 27.68 *μM*. The boundary condition corresponds to per actin filament length of 2π *μm*. Solving this numerically in Python using scipy.integrate, we find that [T-actin]=138.38 *μM*. Please note that while this value is significantly higher than the concentration of actin used in experiments, the value is just one of many degenerate (k_grow_, [T-actin]) value pairs that would numerically satisfy the equations (1) and (2).

To capture essential chemical reactions observed in VASP droplets, filament extension is modeled as a deterministic force-sensitive process as described above along with stochastic Monte Carlo sampling of crosslinker binding and its force-sensitive unbinding reactions. Arp2/3 binding, nucleation, and unbinding are all modeled as stochastic reactions.

### Representation of actin filaments

Actin filaments are represented as inextensible fibers represented as a series of linear segments. The upper limit of segment length is 100 nm. CytoSim computes bending energy of the fiber based on the flexural rigidity specified in the input parameters.

### Representation of VASP tetramer

We model VASP as a spherical crosslinker of radius 30 nm with four F-actin binding sites distributed across the surface of the sphere. Cytosim also requires specification of a binding distance parameter between VASP and actin. As we are interested in the changes to ring-shaped actin networks, we used the parameters from our previous study (https://doi.org/10.1101/2023.05.26.542517) that resulted in the formation of ring-shaped networks.

### Representation of Arp2/3

Arp2/3 molecules are modeled as diffusing point particles. The binding, nucleation, and unbinding reactions are considered as shown in Supplemental figure S2f. In our simulations, Arp2/3 nucleation produces filaments of 5 nm in length at a 70° angle with respect to the parent filament.

### Position evolution

The diffusion of VASP, points along the actin filaments and Arp2/3 molecules are considered in Cytosim through a Langevin equation framework. For a particle i, the coordinates are given by xi = {xi1, xi2, xi3}. The position along each of the dimensions j is evolved according to the stochastic differential equation given by,

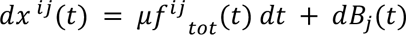

Here, μ is the viscosity of solvent, and *f^ij^_tot_*(*t*) represents the total force acting on the particle at time t. The diffusion term (noise) is given by a random variable sampled from a distribution with mean 0 and standard deviation 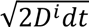, where the diffusion coefficient is given by *D^i^* = *μk_B_T* where T is temperature and k_B_ is Boltzmann constant. Please refer to Supplementary Table S3 for a detailed description of parameters used in this study.

### Analysis of trajectories

Five trajectories were generated for each [Arp2/3] concentration studied. We wanted to understand the distribution of actin within the droplet. Toward this, we discretized the droplet into subvolume shells of varying radii of a finite thickness (δ=25 nm). We calculated the ratio of the local density within shells and the bulk density as,

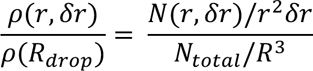

Here, N total refers to the total number of actin monomers in the system. *N*(*r*, *δr*) represents the number of actin monomers in shell of radius r and thickness *δr*.

Further, we also calculated the relative actin enrichment (R.A.E) in the core of the droplet using the following equation.

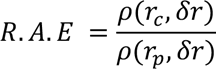

where *r_c_* represents the radius a shell in the core of the droplet and *r_p_* represents the radius of a shell in the periphery of the droplet. We chose *r_c_* = 0.25 *μm* and *r_p_* = 0.9 *μm* in this study.

## Notes

### Competing Interest Statement

The authors have declared no competing interest.

